# Application of environmental DNA-based assessment for upstream-downstream comparison of river macroinvertebrates in a metal-contaminated river

**DOI:** 10.1101/2025.04.02.646737

**Authors:** Noriko Uchida, Yuichi Iwasaki, Ryoichi Kuranishi, Natsuko Ito Kondo

**Affiliations:** Tohoku University; National Institute of Advanced Industrial Science and Technology (AIST); Faculty of Engineering, Kanagawa Institute of Technology; National Institute for Environmental Studies

## Abstract

Environmental DNA (eDNA) is a powerful tool for biological monitoring with the potential to overcome weaknesses of conventional macroinvertebrate surveys in running waters. However, the ability of eDNA to detect changes of macroinvertebrate communities immediately downstream of a perturbation, particularly in upstream-downstream comparisons, has not been adequately explored. To address this issue, we investigated the potential of eDNA-based assessment of a perturbation by comparing the results with macroinvertebrate surveys in a river influenced by the inflow of a metal-contaminated tributary. Both methods revealed distinctly lower richness of taxa and zero-radius operational taxonomic units (ZOTUs) at the metal-contaminated tributary site compared to other study sites. Results from collection of macroinvertebrates indicated that most richness and abundance metrics were significantly reduced at three metal-contaminated sites located 150–750 m downstream from the inflow of the tributary. In contrast, the eDNA-based assessment revealed similar ZOTU richness at a reference site and the downstream, contaminated sites. Although statistically not significant because sample sizes were small, eDNA-based non-metric multidimensional scaling revealed some separation between the reference site and two downstream sites. However, no separation was apparent between the reference site and the site immediately downstream. This result suggested that eDNA at a site 150 m downstream from the inflow was likely affected by downstream drift of eDNA from the upstream reference area. That drift complicated assessment of the community a short distance from the perturbation. The site separation detected by eDNA-based assessment was promising, but the ZOTUs that contributed to the separation were mainly from dipteran taxa rather than from metal-sensitive mayflies, which were significantly lower in abundance at the downstream, contaminated sites. Developing reliable local DNA barcoding information, particularly for these mayflies, may help overcome the limitations of making evaluations over relatively small spatial scales, such as upstream-downstream comparisons.

## 1 Introduction

Freshwater ecosystems provide essential services, including material, non-material, and regulatory functions, that fulfill myriad human needs (Lynch et al., 2023). However, the increasing degradation of these ecosystems and the consequent loss of biodiversity due to several factors including land use change and chemical pollution underscore the urgent need for effective conservation strategies (Persson et al., 2022; Reid et al., 2019; Tickner et al., 2020). Biomonitoring to assess the biological and ecological health of freshwater systems as well as the effectiveness of conservation measures is necessary to ensure nature-positive outcomes.

In freshwater ecosystems such as streams and rivers, benthic macroinvertebrates have frequently been used as bioindicators worldwide (Birk et al., 2012; Buss et al., 2015; Eriksen et al., 2021; Namba et al., 2020). Their sedentary nature, ease of sampling, and diverse sensitivities to environmental stressors make benthic macroinvertebrates particularly valuable for biomonitoring. However, their sorting and taxonomic identification are labor intensive and require specialist expertise in examining their morphological characteristics under a microscope.

Environmental DNA (eDNA) techniques are a promising tool for addressing these challenges faced by traditional biomonitoring methods because they allow for the molecular detection of benthic macroinvertebrate species directly from environmental media such as water and with lower sampling effort (Baird & Hajibabaei, 2012; Blackman et al., 2019; Brantschen et al., 2021; Mächler et al., 2014; Takenaka et al., 2023; Uchida et al., 2020). However, before eDNA can be used as an indicator in environmental assessments, certain aspects of its dynamics need to be elucidated. In particular, the downstream transport and dispersion of eDNA in a river, even without substantial dilution, can complicate its ability to accurately reflect local biotic communities (Deiner & Altermatt, 2014; Jane et al., 2015; Sansom & Sassoubre, 2017). Downstream transport of eDNA has been reported to extend from several hundred meters to over 10 km (Deiner & Altermatt, 2014; Jane et al., 2015; Nukazawa et al., 2018; Sansom & Sassoubre, 2017). Upstream-downstream comparisons at effluent discharge points based on direct collections of benthic macroinvertebrates using traditional biomonitoring methods have been used for environmental impact assessment and bioassessment. The applicability of eDNA to such studies has not been tested. It is of particular importance to determine whether eDNA can capture impacts immediately downstream without being influenced by eDNA from upstream reference areas.

The goal of this study was to examine the potential for assessment of impacts based on eDNA by comparing the results of eDNA analysis with those of macroinvertebrate sampling in a river influenced by a tributary heavily contaminated with metals. For this purpose, we selected study sites where the inflow of the tributary was known to drastically alter downstream benthic macroinvertebrate communities (Iwasaki et al., 2023). This selection allowed us to determine to what extent the influence of upstream communities would compromise the ability of eDNA-based assessment to discern changes in the macroinvertebrate fauna at varying distances downstream from the inflow of the metal-contaminated tributary.

## 2 Materials and Methods

### 2.1 Study Site

We selected a reference (upstream) site (A1) and four metal-contaminated sites (A0, A2–A4) in a tributary (Yunosawa River) of the Waga River in the basin of the Kitakami River, Japan (Figure 1). Site A0, with a catchment area of 4.6 km^2^, was located in a tributary of the Yunosawa River, which receives mine discharge from a closed mine (Akaishi mine). The inflow of the metal-contaminated tributary into the Yunosawa River has been reported to cause increases of the downstream concentrations of metals such as copper and zinc that lead to remarkable reductions of the richness and abundance of benthic macroinvertebrate communities (Iwasaki et al., 2023). To determine whether the influences of eDNA that originated from benthic macroinvertebrates in the upstream reference area on eDNA-based assessments varied with distance downstream, accessible study sites were selected approximately 150 m downstream (Site A2), 600 m downstream (Site A3), and 1350 m downstream (Site A4) from the inflow of the metal-contaminated tributary. Because there was minimal variation in the catchment areas of these three sites (i.e., 19–20 km^2^, based on Yamazaki et al., 2020), we were able to specifically examine the attenuation of the influence of eDNA with downstream distance without considering the influence of dilution by increased river flow. Field sampling, as detailed below, was conducted at these study sites on May 26–27, 2022.

**FIGURE 1.**
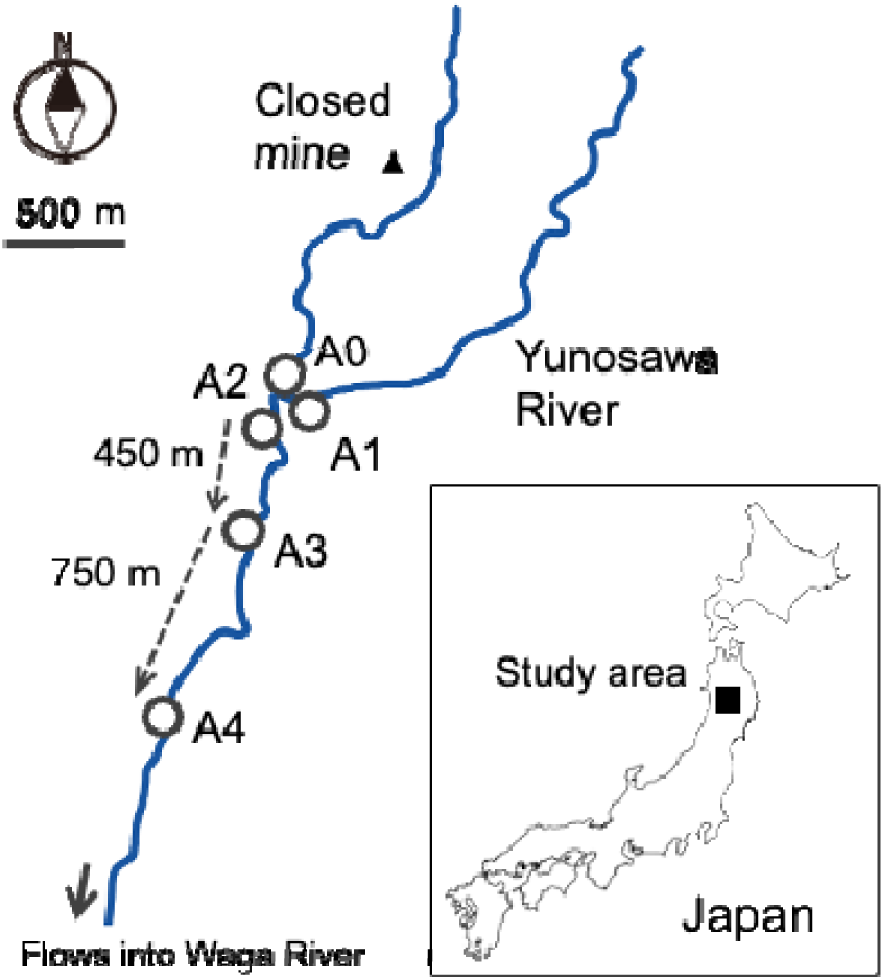
Study sites along the Yunosawa River (Iwate Prefecture, Japan) and distances between them.

### 2.2 Macroinvertebrate Collection and Analysis

Macroinvertebrates were collected at three locations within cobble-dominated riffles at each study site using a 25 cm × 25 cm Surber net (mesh size, 0.355 mm). We followed the quantitative sampling method described in the Basic Survey Manual for the National Census on the River Environment (NCRE) (MLIT, 2016). Macroinvertebrate samples were preserved in jars with 90% ethanol in the field and transported to the laboratory for sorting and identification. For each sample, benthic macroinvertebrates that were sorted using a sieve with a mesh size of 0.5 mm were generally identified to the species or genus level.

To examine differences between benthic macroinvertebrate communities among the study sites based on field sampling, we calculated the taxon richness (number of taxa) and abundance (number of individuals) of three major insect groups (Ephemeroptera, Trichoptera, and Diptera) as well as total taxon richness and total abundance. The taxon richness and abundance of stoneflies (Plecoptera) were not calculated because of their limited richness (≤4 taxa) and abundance (<20 individuals) at these sites. In addition, the abundances of baetid, ephemerid, and heptageniid mayflies (families Baetidae, Ephemeridae, and Heptageniidae, respectively) were calculated. These three mayfly families have contrasting sensitivities to metals (Iwasaki et al., 2018). Reduced abundances of metal-sensitive ephemerid and heptageniid mayflies have been observed in metal-contaminated rivers, but baetid mayflies can still be found in rivers heavily contaminated with metals.

### 2.3 Water Quality Measurements

We generally followed the protocols for water quality measurements described in our previous study (Iwasaki et al., 2023). Measurements of pH and electrical conductivity were performed on-site using portable water quality meters (LAQUAtwin pH-22B and EC-33B, Horiba, Kyoto, Japan). Grab samples of water for dissolved trace metals (Cu, Zn, Cd, and Pb) and major ions (Na, K, Ca, Mg, Cl, and SO_4_) were collected by filtering the water through a hydrophilic PTFE filter (pore size, 0.45 μm) and were immediately refrigerated in the field. Ultrapure nitric acid was added to water samples for trace metal analysis in the laboratory, and concentrations of dissolved Cu, Zn, Cd, and Pb were determined using an inductively coupled plasma mass spectrometer (Element XR, Thermo Fisher Scientific, MA, US), in accord with U.S. Environmental Protection Agency (1994). Concentrations of major ions were analyzed using an ion chromatograph (Dionex ICS-1100/2100, Thermo Fisher Scientific).

## 3 Extraction of eDNA, Library Preparation, Bioinformatics

### 3.1 Sampling

Sampling for eDNA was conducted at the same location and on the same days but immediately before macroinvertebrate and water quality surveys. Three replicates were taken at each site from the surface water of the river at the end of a riffle. One liter of water was filtered through a 0.45-um Sterivex filter (Merk Millipore, MA, US) using a disposable, sterile, 50-mL syringe. The filters were finally aerated and drained thoroughly before being injected with 2 mL of RNALater (Thermo Fisher Scientific) and stored at ambient temperature in a plastic bag during field transportation. It was later transferred and stored in a refrigerator at −20°C in the laboratory.

### 3.2 Molecular Analyses

DNA was extracted by adding 720 mL of buffer ATL and 80 µL of 20 mg/mL Proteinase K solution directly to the Sterivex cartridge and incubating the solution at 56°C overnight (Spens et al., 2017). The liquids were removed from the inlet of the cartridge with a luer lock syringe and transferred to a 5-mL LoBind tube. The subsequent extraction was performed via a protocol very similar to that of Spens et al. (2017) using DNeasy Blood and Tissue Kit (Qiagen, Hilden, Germany). The final elution volume of buffer TE was 100 µL, which was stored until further processing at −20°C.

Samples were amplified using a 142-bp fragment of the mitochondrial cytochrome c oxidase subunit I (COI) marker with a universal primer set that targeted macroinvertebrates—especially Ephemeroptera, Plecoptera, Trichoptera, and Diptera—using the forward primer fwhF2 (Vamos et al., 2017) and reverse primer EPTDr2n (Leese et al., 2021). This primer pair included a modification that included the Nextera transposase sequences and shifts for a two-step PCR approach (Elbrecht et al., 2016). The first PCR was conducted in a 10-µL volume with three technical replicates containing 1× Multiplex PCR Master Mix (Qiagen Multiplex PCR Plus Kit, Qiagen) and 0.2 µM of each primer. The volume was then brought up to 10 µL with Nuclease-free water and 2 µL of DNA template. The first PCR, which was performed following Leese et al. (2021), started with initial denaturation at 95°C for 5 min followed by 10 cycles of denaturation at 95°C for 30 s, annealing at 64–54°C for 90 s (touch-down PCR), and extension at 72°C for 30 s. Thirty cycles of PCR were subsequently performed with initial denaturation at 95°C for 30 s, annealing at 54°C for 90 s, and extension at 72°C at 30 s. The final extension was performed at 68°C for 10 min, and the plates were cooled to 4°C. Three technical replicates of the first PCR products were pooled and purified using AMPure XP (Beckman Coulter, Bread, CA, USA). The second PCR mixtures were prepared with 1× Multiplex PCR Master Mix, 0.2 µM of each primer of Nextera XT index with Illumina adaptor sequences, and 2 µL of the purified first PCR product. The volume was brought up to 10 µL with Nuclease-free water. The PCR started with initial denaturation at 95°C for 15 min followed by 10 cycles of denaturation at 95°C for 30 s, annealing at 64°C for 90 s, and extension at 72°C at 60 s. The final extension was performed at 72°C for 5 min, and the plates were cooled to 4°C. The amplicons were purified in the same manner as the first PCR products, but with a size selection to remove remaining primers and primer dimers (0.8× SPRIselect; Beckman Coulter). The concentrations of PCR products were quantified with the Qubit dsDNA High Sensitivity Kit (Thermo Fisher Scientific) and adjusted and pooled so that each sample had an equal concentration. The final amplicon length of the pooled sample was measured with Tape Station D1000 Screen Tape (Agilent, Santa Clara, CA, USA). The paired-end sequencing was performed using MiSeq (Illumina) with Reagent Kit v2 (300 cycles) with a 10% PhiX spike in accordance with the manufacturer’s instructions at the Genome Diversity Centre, ETHZurich, Switzerland.

### 3.3 Bioinformatics

The raw sequence data were first checked using FastQC (Andrews, 2010). Whole bioinformatics were performed under the environment of USEARCH v.11.0.667 (Edgar, 2010). The PhiX, low-complexity reads, and primer sequences (fwhF2/EPTDr2n) were removed. The forward and reverse reads were then trimmed to 100–300 nucleotides to remove low-quality bases. Chimeric sequences were removed. Reads with ambiguities or a maximum expected error greater than 2 were removed. The demultiplexed forward and reverse reads were merged and clustered using UNOISE3 (USEARCH v.10.0.240) to make zero-radius OTUs (ZOTUs) with an additional clustering at 99% identity of ZOTUs. The ZOTUs were inferred based on error rate model estimates, and the paired reads were merged into one sequence with a minimum overlap of 30 bases. Finally, ZOTUs that were found in relative proportions >0.1% in one of the negative controls were also discarded from all the samples. ZOTU sequences were assigned taxonomically with the SINTAX algorithm (Edgar, 2016) implemented in the USEARCH package using a confidence threshold of 80% for a family level and 70% for a genus level, and the MIDORI2 reference database (Leray et al., 2022).

### 3.4 Statistical Analyses

For macroinvertebrate metrics based on benthic macroinvertebrate samples and the richness of ZOTU based on eDNA analysis, differences between metal-contaminated sites (A0, A2–4) and the reference site (A1) were examined using Dunnett’s multiple comparison test following a one-way analysis of variance (ANOVA). The richness and abundance metrics based on benthic macroinvertebrate samples were log_10_-transformed to align with the assumptions of this analysis.

For benthic samples, nonmetric multidimensional scaling (NMDS) based on Bray-Curtis dissimilarities (Manly & Navarro Alberto, 2016) was performed to compare macroinvertebrate communities among the study sites. Square-root-transformed abundances of macroinvertebrate taxa were used to calculate the Bray-Curtis dissimilarities. Similarly, for eDNA, NMDS based on Jaccard dissimilarities using presence-absence data of ZOTUs was performed. The NMDS analysis was performed using the R package “vegan” (v.2.6.4; Oksanen et al., 2019) in R (v.4.3.1; R Core Team, 2023). For eDNA, similarity percentage analyses (SIMPER, Clarke, 1993) were conducted to identify ZOTUs that contributed to the separation observed among sites.

## 4 Results and Discussion

### 4.1 Water Quality and Macroinvertebrates

The increase of downstream metal concentrations and significant impacts on many macroinvertebrate metrics (Figure 2 and Table S1) associated with the inflow from the heavily metal-contaminated tributary were consistent with findings reported by Iwasaki et al. (2023). While metal concentrations at the reference site (A1) were well below the US EPA water quality criteria at a water hardness of 15 mg CaCO_3_/L (Cu: 1.8 μg/L; Zn: 24 μg/L; Cd: 0.17 μg/L; Pb: 0.30 μg/L), concentrations of metals, with the exception of Pb, at the metal-contaminated sites (A0, A2–A4) exceeded these criteria. Particularly notable were the exceedances of the Cu criterion; concentrations at A0 and A2 were approximately 50 and 16 times, respectively, the criterion value. This result indicates that there were significant ecological risks at these contaminated sites.

**FIGURE 2.**
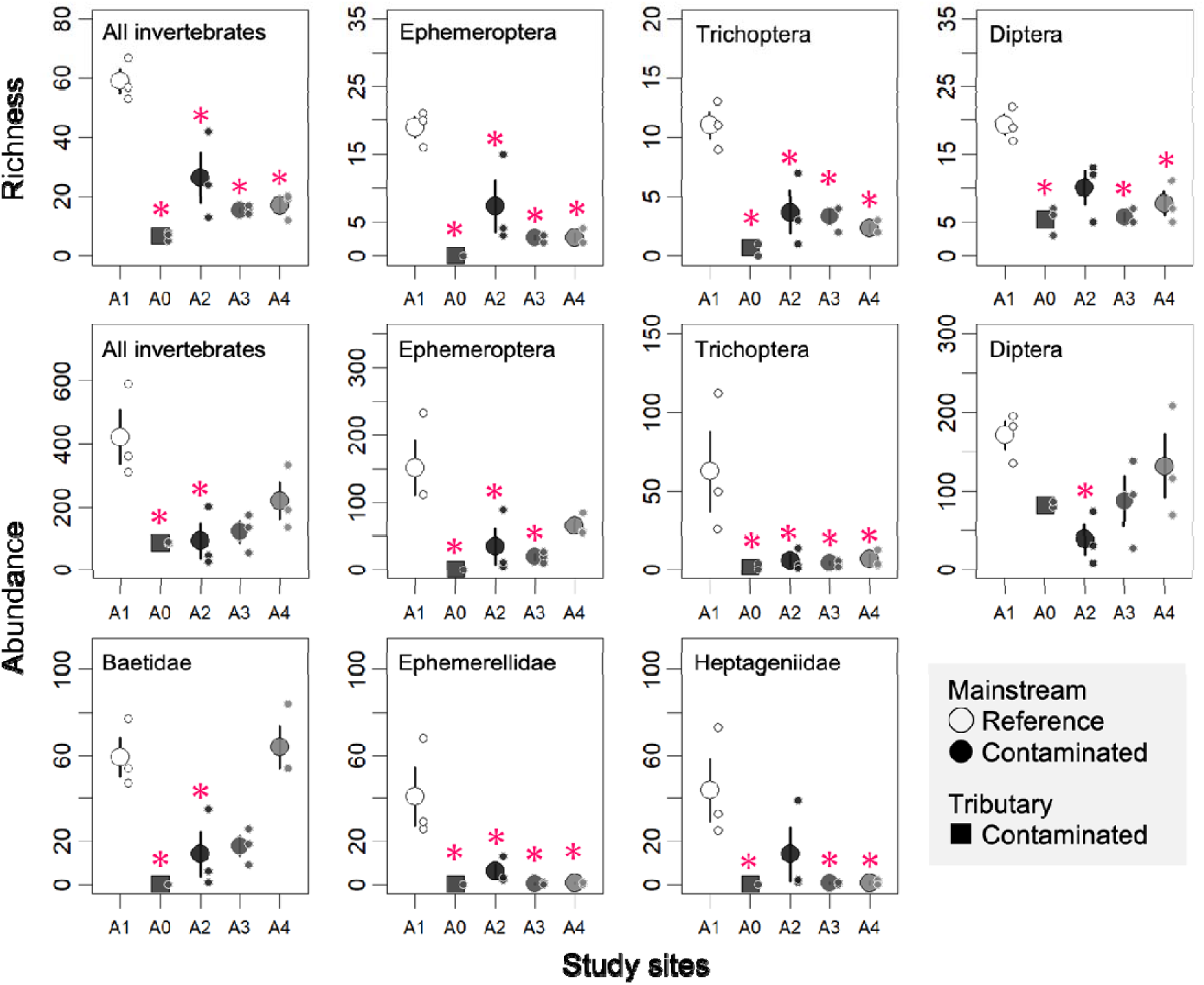
Richness and abundances of all macroinvertebrates and three dominant orders (Ephemeroptera, Trichoptera, and Diptera), and abundances of three dominant families (Baetidae, Ephemerellidae, and Heptageniidae) at the study sites. Large symbols and error bars represent means and ±1 standard error, respectively. Small symbols indicate individual observed values. Asterisks indicate significant differences (*p* < 0.05, Dunnett’s multiple comparison) compared to the reference site (A1).

As predicted from the metal concentrations, ∼90% of the macroinvertebrates collected at A0 were chironomids (Diptera), and no mayflies were collected (Figure 2). While certain macroinvertebrate metrics, such as the abundance of baetid mayflies (Baetidae), that are often found in metal-contaminated rivers, were not significantly different from those at the reference site (A1), the majority of abundance and richness metrics were significantly lower at the downstream, metal-contaminated sites (A2–A4) than at A1 (Figure 2). Despite the limited statistical power due to the small sample size (three replicates per site), these results revealed a clear distinction between macroinvertebrate communities at A1 and those at other contaminated sites (A0, A2–A4). The NMDS result based on Bray-Curtis dissimilarities among all macroinvertebrate samples collected at all study sites (Figure 3a) showed that communities at the tributary site (A0) were clearly separated from those at the reference site (A1) and the contaminated sites (A3 and A4), while those at site A2 plotted between these four sites. The taxon richness of one of the macroinvertebrate samples collected from A2 was comparable to that at the reference site (A1), and the variations of the richness and abundance metrics were larger at A2 than at the other sites (Figure 2). These results at A2 may have been due to a temporary occupation by macroinvertebrate taxa that drifted from the reference area immediately upstream, including site A1. The importance of such drift in assessing the impact of metal contamination on macroinvertebrates has been discussed in previous studies (Beltman et al., 1999; Maret et al., 2003).

**FIGURE 3.**
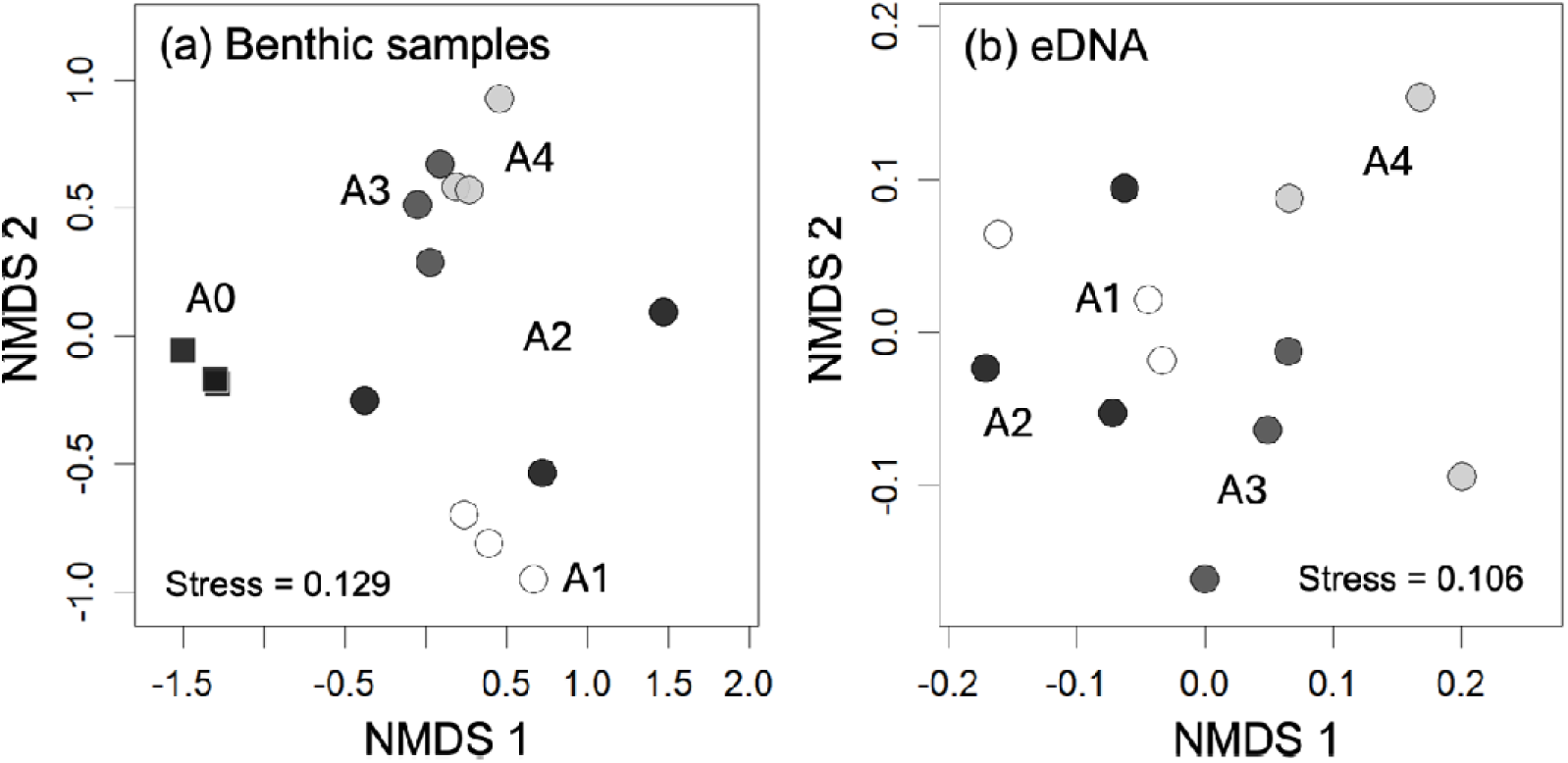
Results of nonmetric multidimensional scaling (NMDS) based on (a) Bray-Curtis dissimilarities for benthic samples and (b) Jaccard dissimilarities for eDNA samples. Individual data points are the three replicates at each study site. Dashed lines have been added to identify study sites more clearly. For the eDNA samples, we excluded A0 to clearly reveal differences among A1–A4 (see Figure S1 for the result including A0).

### 4.2 Analysis of eDNA

A total of 2,892,569 raw sequence reads were obtained from all samples, and 2,793,679 of them passed the sequence quality filter and were used to create ZOTUs (average 186,245 reads/sample). A total of 3,903 ZOTUs were formed, and 838 of them were matched with the threshold and were identified as Insecta. All six ZOTU richness metrics at A0 (i.e., Total ZOTUs, EPTD, Ephemeroptera, Plecoptera, Trichoptera, and Diptera) were 24–74% lower than those at the reference site, A1 (Figure 4, *p* < 0.05). However, in contrast to the macroinvertebrate results, the ZOTU richness metrics were similar at the reference site and downstream, contaminated sites. Indeed, despite the significant reductions observed in the macroinvertebrate collection, ZOTUs of Baetidae, Ephemerellidae, and Heptageniidae detected at A1 were, in most cases, also detected at the downstream, contaminated sites with similar read numbers (Table S2). These results clearly indicated that the ZOTU richness metrics using eDNA failed to detect the significant reductions of macroinvertebrates observed at the downstream, contaminated sites (i.e., A2–A4).

**FIGURE 4.**
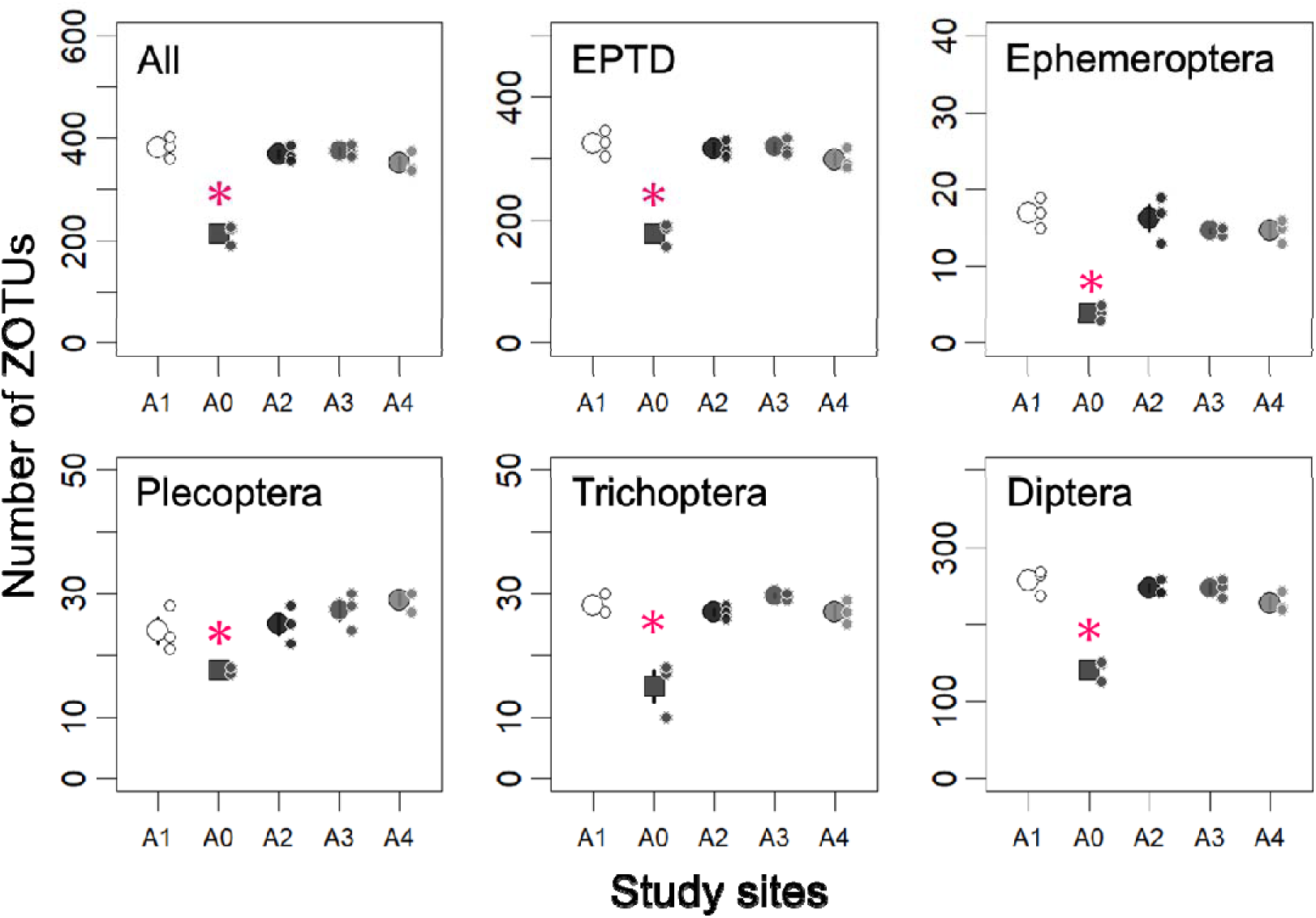
Number of zero-radius operational taxonomic units (ZOTUs) for all taxa (including Arthropoda, Annelida, and Discosea), EPTD (Ephemeroptera, Plecoptera, Trichoptera, and Diptera), Ephemeroptera, Plecoptera, Trichoptera, and Diptera. Large symbols and error bars represent means and ±1 standard deviation, respectively. Small symbols indicate individual raw observed values. Asterisks indicate significant differences (*p* < 0.05, Dunnett’s multiple comparison) compared to the reference site (A1).

The NMDS analysis of eDNA data indicated that community dissimilarities between A0 and other sites were remarkably large (Figure S1). Although this result was consistent with the results based on the ZOTU richness metrics, it made detailed comparisons between A1 and the downstream, contaminated sites difficult. The NMDS result based on eDNA data at the mainstream sites (i.e., except A0; see Figure 3b) visually revealed a modest separation between A1 and A2, and between A3 and A4. However, the difference between even A1 and A3 or A4 was not statistically significant (*p* > 0.05; permutation tests with pseudo-F ratios with Bonferroni adjustment using the “adonis2” function in R package “vegan”), most likely because of the small sample size (n = 3).

Selection of ZOTUs that contributed significantly to the separation between A1 and A2 as well as between A3 and A4 by SIMPER analysis most often resulted in ZOTUs from Diptera (13 of the selected 30 ZOTUs), followed by 1–6 ZOTUs from six orders (Table S3): Coleoptera (6), Hemiptera (4), Lepidoptera (3), Plecoptera (3), Trichoptera (2), and Neuroptera (1). No ZOTUs from Ephemeroptera were selected as “indicator” ZOTUs, likely because of the constant detection of ZOTUs from Baetidae, Ephemerellidae, and Heptageniidae (Table S2), which were rarely collected in the benthic samples at the contaminated sites (A2–A4). These results based on eDNA detection were rather counterintuitive given that ephemerellid and heptageniid mayflies have been reported to be taxa sensitive to metals (Iwasaki et al., 2018). The occurrences of many of those selected ZOTUs were typically only at A1 and A2 or at A3 and A4. However, because of the lack of information about the metal tolerance of selected “indicator” taxa detected by eDNA, it is difficult to say whether metal contamination was the cause of these characteristic occurrences. For example, some chironomids have been reported to be metal-sensitive (Iwasaki et al., 2009), but a rigorous comparison of eDNA and benthic collection results is difficult to perform, largely because of the challenges of morphological identification at the species level.

Despite the possibility that the responses of the selected ZOTUs may have been caused by factors other than metal contamination (e.g., natural longitudinal variations), it is encouraging that the NMDS analysis based on eDNA data made it possible to distinguish between the reference site (A1) and the contaminated sites (A3 and A4), although the results were not statistically significant. In contrast, eDNA results obtained at A2, ∼150 m downstream from A1, were likely to have been substantially affected by the downstream drift of eDNA originating from communities in the reference area, including A1 (Figure 3b). This likelihood suggests the difficulty of assessing changes in macroinvertebrate communities using eDNA at such short distances, no matter how large an impact is observed at a downstream site. The use of macroinvertebrate eDNA as an alternative to conventional upstream-downstream comparisons of benthic samples is thus not yet straightforward. A promising solution to this problem would be to use environmental RNA (eRNA), which is more rapidly degraded in the field than DNA and therefore reflects local and immediate biological signals (Jo et al., 2022; Marshall et al., 2021; Miyata et al., 2022).

In the present study, dipteran taxa were often used as indicator ZOTUs. We found that it was necessary to increase the number of sequence reads for the targeted species and to reduce the amount of “not assigned” data in the process of taxonomic assignment in order to obtain robust eDNA data for assessment. In the present study, the number of ZOTUs ultimately used for analysis differed across taxonomic groups. Specifically, the family Chironomidae within the order Diptera included 269 ZOTUs that were retained out of the total of 838 ZOTUs. However, we retained only 26 ZOTUs in the order Ephemeroptera out of the total of 838 ZOTUs. One possible reason for this difference was that the COI primers used in this study amplified dipteran DNA more effectively. This primer set has the advantage for detecting freshwater macroinvertebrates (Ephemeroptera, Plecoptera, Trichoptera, and Diptera) because it minimizes the amplification of non-target DNA, such as DNA from algae (Leese et al., 2021; Mächler et al., 2021). However, for purposes of minimizing intraspecific amplification biases, primers targeting the phylogenetically conserved mitochondrial 16S rRNA region (Takenaka et al., 2023, 2024) may be more suitable. Another reason may be that the disparity in the number of sequences registered as reference DNA across species can affect the taxonomic identification process. Indeed, the number of registered sequences in the database for Diptera (1,244,578) is substantially greater than that for Ephemeroptera (23,945). In particular, the database for Japanese Chironomidae (a family within Diptera) is well established and includes a large number of registered sequences (Chironomid DNA Barcode Database; https://www.nies.go.jp/yusurika/en/contents/methods.html). Given the high genetic diversity of insects (Tojo et al., 2017) and the inherent concerns about the accuracy of existing DNA reference databases (Keck et al., 2023), it is crucial that reliable local DNA barcoding information be expanded (Mugnai et al., 2023). Such developments could help to overcome the limitations associated with evaluations at relatively small spatial scales, such as the upstream-downstream comparisons made in this study, and could ultimately contribute to informed consideration of the optimal uses of the macroinvertebrate eDNA approach.

**Table S1.** Water quality at study sites. Limits of quantification for Cu, Zn, Cd, and Pb were 0.1, 0.5, 0.0001, and 0.025 μg/L, respectively. US EPA water quality criteria at a water hardness of 15 mg/L Cu, Zn, Cd, and Pb are 1.8, 24, 0.17, 0.30 μg/L, respectively.

| Site | Unit | A0 | A1 | A2 | A3 | A4 |
| --- | --- | --- | --- | --- | --- | --- |
| pH |  | 7.7 | 7.8 | 8.1 | 8.3 | 8.0 |
| EC | µs/cm | 56 | 69 | 66 | 64 | 69 |
| Cu | µg/L | 90.9 | 0.2 | 28.7 | 20.9 | 17.5 |
| Zn | µg/L | 243.8 | <0.5 | 76.0 | 44.9 | 36.4 |
| Cd | µg/L | 5.55 | <0.001 | 1.73 | 1.07 | 0.92 |
| Pb | µg/L | 0.11 | <0.025 | 0.03 | <0.025 | 0.05 |
| Ca | mg/L | 3.1 | 3.8 | 3.6 | 3.6 | 3.7 |
| Mg | mg/L | 2.2 | 2.1 | 2.1 | 2.1 | 2.0 |
| Na | mg/L | 5.5 | 6.5 | 6.4 | 6.5 | 6.7 |
| K | mg/L | 1.0 | 0.9 | 0.9 | 0.9 | 0.9 |
| Cl | mg/L | 6.7 | 6.3 | 6.4 | 6.5 | 6.4 |
| SO<sub>4</sub> | mg/L | 13.4 | 4.0 | 6.9 | 6.2 | 5.8 |
| Hardness | CaCO<sub>3</sub> mg/L | 16.9 | 18.3 | 17.6 | 17.6 | 17.4 |

**Table S2.**
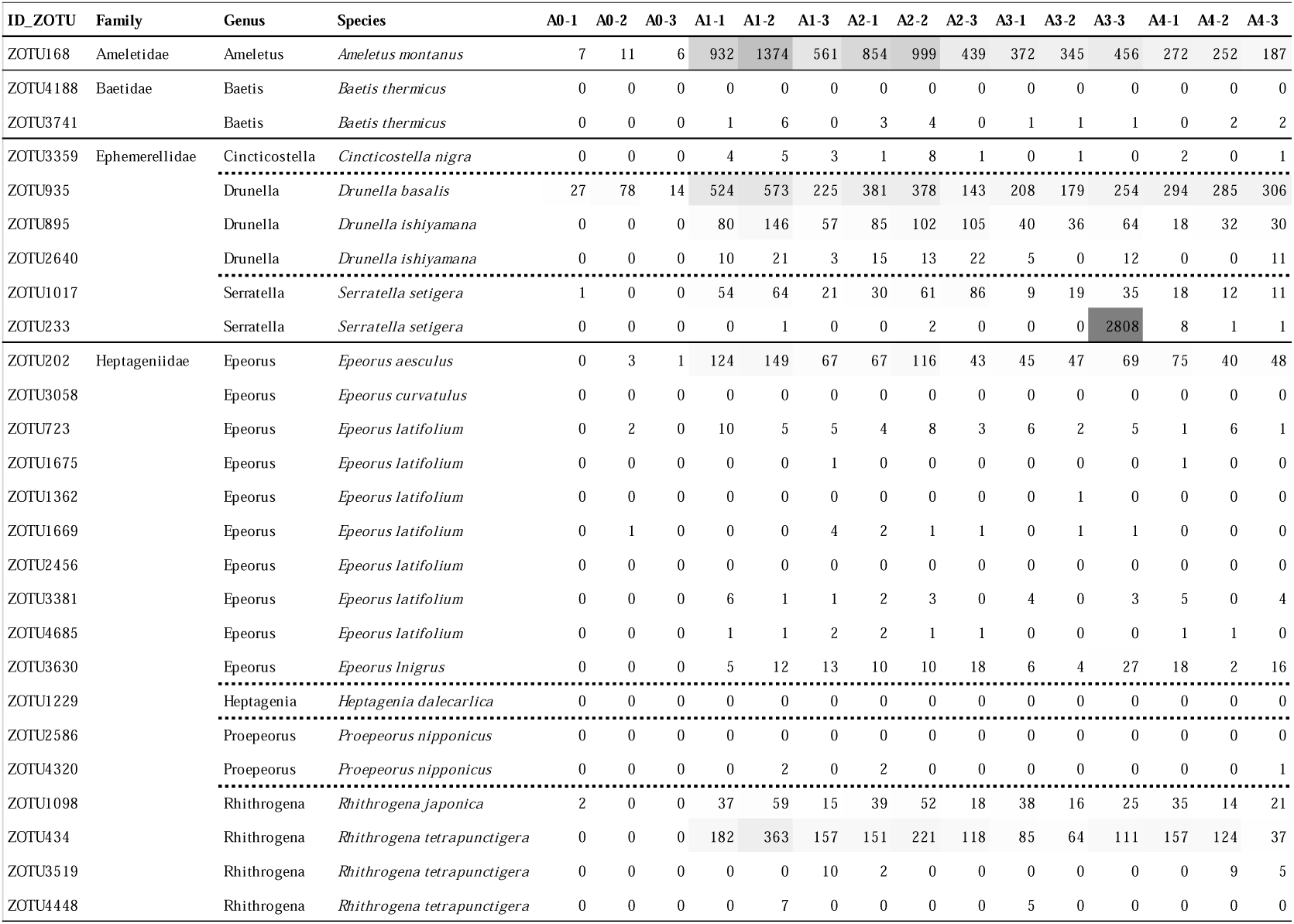
Number of reads of Ephemeropteran ZOTUs with taxonomic assignment results.

**Table S3.**
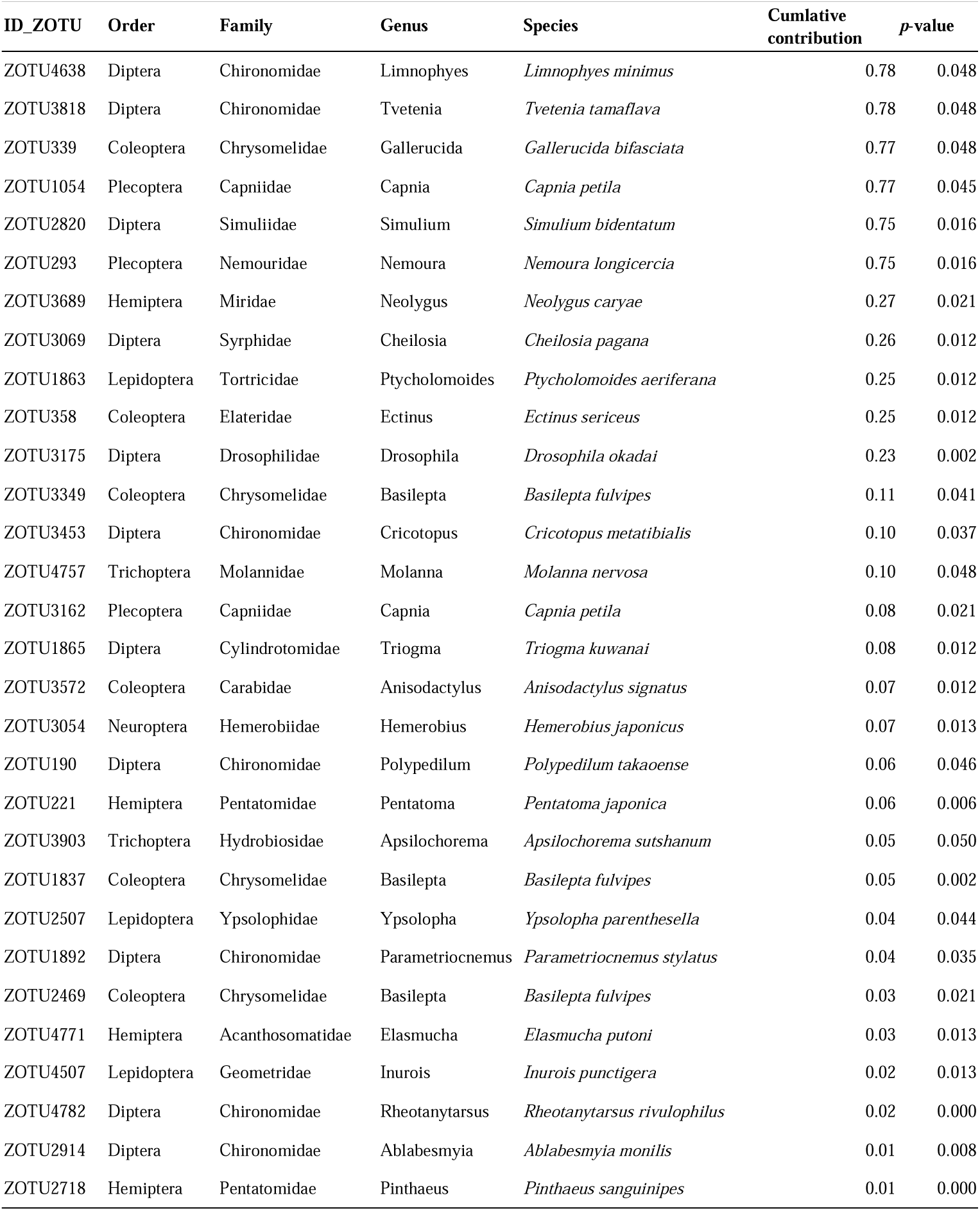
Results of SIMPER analysis ordered by cumulative contribution with *p* < 0.05.

**FIGURE S1.**
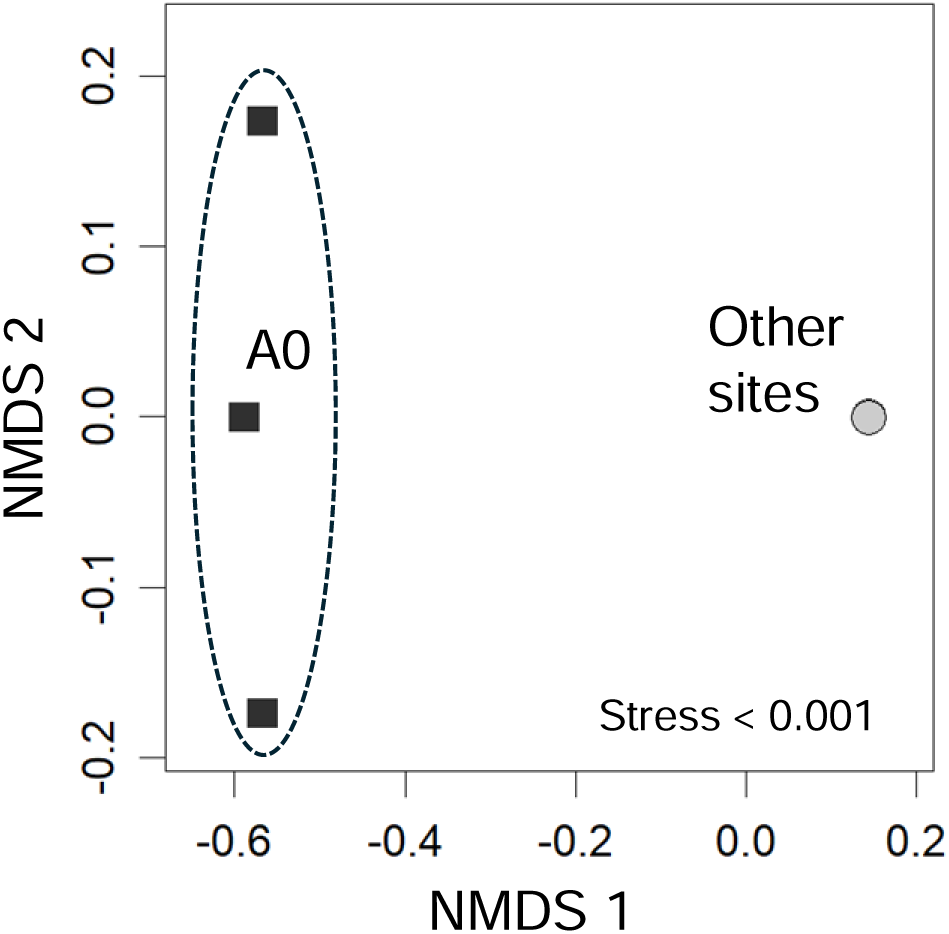
Results of nonmetric multidimensional scaling (NMDS) based on Jaccard dissimilarities for eDNA samples. Symbols for sites A1–A4 were located in almost the same positions in this figure and could not be distinguished (see Figure 3b).

